# Inferring single-cell heterogeneity of bacteriophage lysis-associated life-history traits from population-scale dynamics

**DOI:** 10.1101/2025.03.25.645349

**Authors:** Marian Dominguez-Mirazo, Ran Tahan, Shay Kirzner, Debbie Lindell, Joshua S. Weitz

## Abstract

Phage-induced lysis of bacteria transforms population dynamics, community structure, and ecosystem functioning. Scaling up infected cell fate to quantify population- and ecosystem-scale impacts requires estimates of viral life history traits, including underlying heterogeneity in the timing, efficiency, and outcome of lytic infections. However, the variability of lysis-associated phage traits remains poorly characterized. Here, we infer single-cell heterogeneity in lysis-associated traits for an ecologically relevant system: Syn9, a T4-like cyanophage infecting *Synechococcus* strain WH8109, a representative of globally abundant marine cyanobacteria. We estimate the heterogeneous distribution of latent period and burst size using a nonlinear model of infection dynamics applied to population-scale time series data. We then validate our inference approach using a single-cell assay – demonstrating the feasibility of inferring phage trait heterogeneity from population data even in the absence of single-cell experiments. The variation in Syn9’s latent period exceeds that previously found in coliphages and reinforces the limitations of representing viral traits with a single value. Moreover, by partitioning lytic events via the inferred heterogeneous latent period distribution, we show that realized burst size variability is largely explained by differences in latent period, providing a path forward to measure and integrate trait (co)variation into population and ecosystem models.

## 1 Introduction

Bacteriophage (phage) adsorb, infect, and lyse sensitive bacteria, transforming the fate of cells and populations. Cumulatively, viruses of microbes are estimated to kill a substantial fraction of marine microbes daily in surface waters [1] (with estimates spanning 1-5% in oligotrophic gyres [2, 3] to ∼10-30% in transition zones [3, 4]), and potentially higher levels during blooms [5–7]. The lysis of marine microbes leads to the release of organic matter back into the microbial loop [7–10] and may generate sticky aggregates that can lead to increases in export of organic matter in the water column [6, 11, 12].

The viral lytic life cycle includes multiple stages: phage adsorb to bacterial cell surfaces, inject their genetic material into the cytoplasm, hijack cell machinery to produce new infectious virions, and finally – if the infection is successful – lyse the host cell, releasing viral progeny back into the environment. The quantitative features of different stages can be represented in terms of ‘viral life-history traits’, including the adsorption rate, latent period (time between attachment and lysis) and the burst size. Conventionally, these viral life history traits are quantified via a single number estimated from a population, i.e., a value of that trait in a population that is presumed to be representative of trait values in the population as a whole. Quantitative estimates of lysis-associated life-history traits have been shown to differ between phage-bacteria pairs [13–15] and can vary with environmental conditions (e.g., temperature [13, 16]) or nutrient availability (e.g., nitrogen and phosphorus [17, 18]).

Phage life history traits can exhibit considerable variability even within a given phage-host pair under fixed environmental conditions. More than 80 years ago, Max Delbrück reported significant variation in burst size for *E. coli* B and phage ‘alpha’ (now known as T1), that ‘cannot be accounted for by variations in the size of the bacteria alone’ [19]. Quantifying the source and extent of heterogeneity is relevant at cellular and ecological scales. Intrinsic variation of lysis-associated phage life-history traits influences microbial population dynamics, e.g., viruses released during early lysis events within a population of infected cells can go on to infect new cells earlier than those viruses that lyse later on [20–24]. Although latent period variability has been characterized within some host-phage systems [25], these studies typically focus on *E. coli* -infecting phage where the latent period is tightly regulated [23, 26–28]. Whether or not such variability is common or conserved across diverse phage-host systems remains unresolved in part due to the technical challenges of single-cell studies.

Cyanobacteria are the most abundant photosynthetic organisms on the planet [29, 30]. Cyanophages play a significant role in shaping the ecology and evolution of cyanobacteria [31–33] and, in turn, influence the global flow of carbon and nutrients [1, 9]. Here, we explore variability in lysis-associated life history traits for Syn9, a lytic T4-like cyanophage infecting *Synechococcus* sp. WH8109, a representative of globally abundant marine cyanobacteria. To do so, we extend a previously developed nonlinear mathematical modeling framework [20] to infer bacteriophage latent period distributions from population-scale time series experiments. By fitting time series data to a lytic infection model that considers latent period variability, we predicted the lysis-associated life-history traits of Syn9. We then adapted a single-cell lysis assay [34] to empirically measure the latent period of Syn9 in individual cells. As we show, population-level data can be used to accurately characterize latent period variation within single cells. Moreover, intrinsic variability in the latent period also explains variation in burst size. Together, this study characterizes latent period and burst size variability and their interrelation in an ecologically relevant system, and offers a time-series inference framework for similar investigations in other phage-bacteria systems even when single-cell measurements are infeasible or unavailable.

## 2 Results

### 2.1 Lysis time distributions recapitulated from population-level time series

One-step growth curves, a common method for estimating bacteriophage life-history traits, rely on measuring the accumulation of free virus during a single round of infection [35]. The latent period is typically reported as a single value, approximated by the time of the first visible burst [35]. However, in the presence of intrinsic variability in the latent period, this estimate of a single trait value is biased – treating early events as representative of a heterogeneous population [20]. One-step growth curves also fail to provide enough information to recapitulate latent period distributions (described in terms of mean and variance). In contrast, prior work hypothesized that time series of multiple rounds of infection – multi-cycle growth curves – potentially allow for the accurate prediction of underlying trait distributions [20].

Here, we set out to test this hypothesis by characterizing the latent period distribution of the Syn9 phage during infection of the marine cyanobacterium *Synechococcus* sp. WH8109 (*Synechococcus* from here on). A ‘multi-cycle growth curve’ was obtained by inoculating phage in a population of *Synechococcus* growing in liquid medium. Plaque assays were used to quantify the number of infective viruses at multiple time points from 0 to 24 hours after inoculation, capturing multiple cycles of infection over a time period consistent with a single, uninfected host division period (Figure 1A). We used a nonlinear, differential equation model to describe the interactions between lytic phage and bacteria (Figure 1B) [20]. The model represents a microbial population in which the timing of infected cell lysis follows an Erlang distribution, a continuous probability distribution that results from the sum of exponentially distributed events (Figure S1). This distribution can be defined with the mean and coefficient of variation (CV) of the latent period (see Materials and Methods). We used a previously described Bayesian inference framework to fit the model to the ‘multi-cycle growth curve’ [20], inferring the life-history trait values, including the latent period distribution, that are compatible with the time-series data (Figure 1C). Single-cell data from this host-virus system was not used to develop the population-level framework nor used as part of population-level inference (see Table S1 for all parameter estimations, and Figure S2 for chain convergence analysis). A conventional point estimate based on the timing of the first burst in a one-step growth assay would place the latent period between 4 and 4.5 hr (Figure 1C). In contrast, the Bayesian inference yields a mean latent period of 7 hr (95% CI = 6.7–7.5 hr) and a predicted coefficient of variation (CV) of 0.15 (95% CI = 0.13–0.18). This implies that individual infected cells are expected to lyse as early as ∼4 hr and as late as ∼10 hr —a 2.5-fold difference between the earliest and latest bursts (Figure 1D). This disagreement between the latent period as inferred via widely used one-step growth assays and that inferred via a mechanistic, population dynamics framework suggests the need to measure latent period distributions one infection at a time via a single-cell assay.

**Figure 1:**
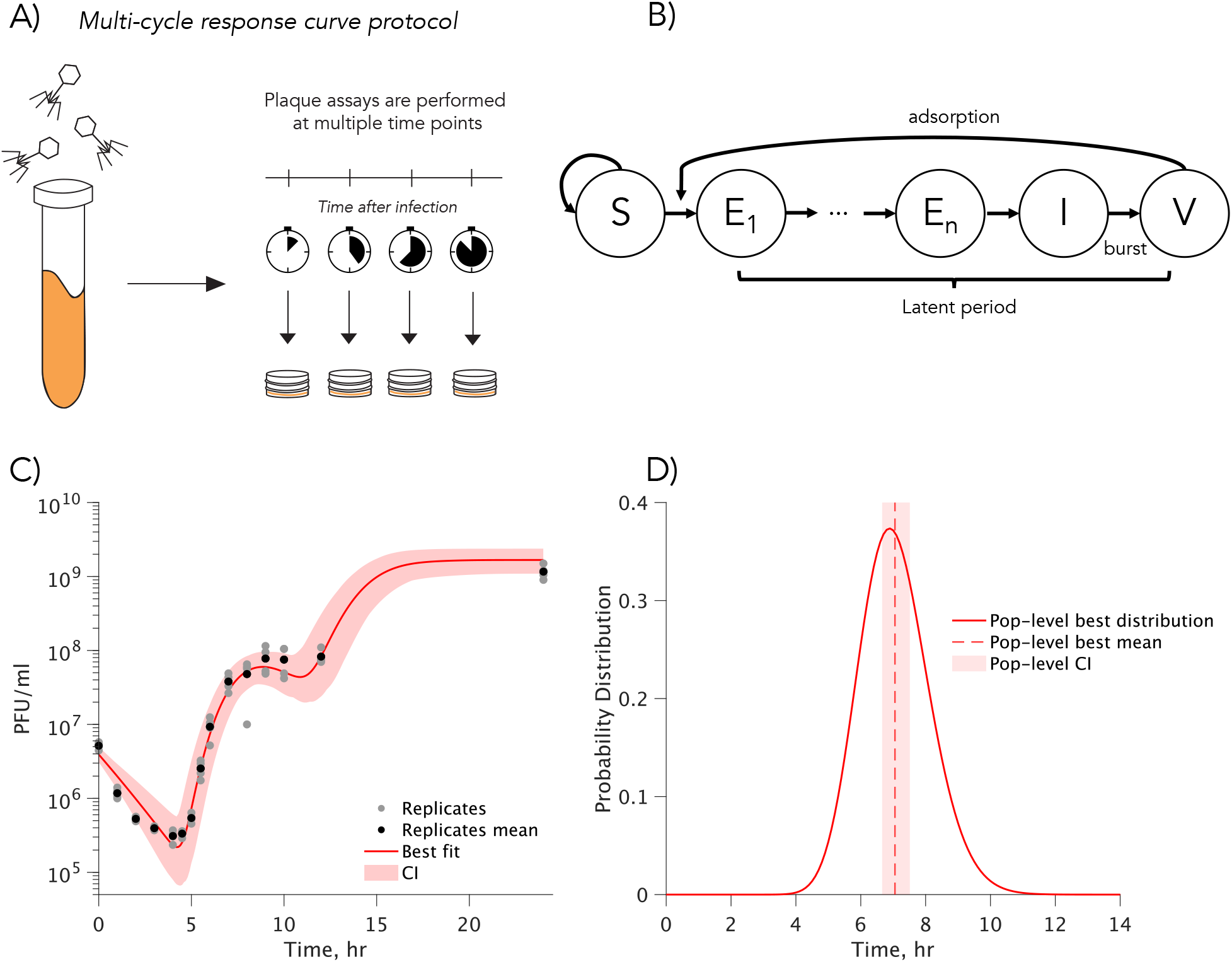
Predicting single-cell heterogeneity from population-level data. We model lysis time heterogeneity in a *Synechococcus* population infected by cyanophage Syn9 using viral density time-series data. **A)** A ‘multi-cycle response curve’ is generated by infecting the cyanobacterial population with the cyanophage. Samples are collected at multiple time points, and viral density is quantified via plaque assays. **B)** In this compartmental model of lytic viral infection, infectious viral particles (*V*) adsorb to susceptible cells (*S*). Infected cells progress through several intermediate stages of infection (*E*_1_ to *E*_*n*_) before reaching the actively infected state (*I*). Infected cells lyse, releasing new viral particles. The mean time for a cell to transition from the first intermediate infection stage (*E*_1_) to final lysis is the mean latent period. The number of intermediate infection stages (*n*) modulates the coefficient of variation (CV) of the latent period distribution. **C)** Using data from four experimental replicates, we determine the parameters that best fit the lysis infection model to the observed viral dynamics from the multi-cycle response curve. Shaded regions represent a 95% confidence interval (CI). **D)** Among the predicted parameters, we estimate the latent period distribution of the population. The mean latent period is predicted to be 7 hr [CI: 6.7, 7.5], and the CV is 0.15 [CI: 0.13, 0.18]. Table S1 lists all predicted parameter values.

### 2.2 Single-cell heterogeneity in lysis time

We set out to test population-level predictions of latent period heterogeneity that is not reflected in conventional inference of one-step growth curves. We devised a protocol for single-cell detection of lysis time adapted from previous work that determines the variability in single-cell virus production [34]. Briefly, we infected a *Synechococcus* population growing in liquid media with phage Syn9 at a relatively high host density and phage-host ratio to maximize contacts (see Methods). After 15 minutes, the population was diluted to reduce phage-bacteria encounters and ensure that all infections resulted from encounters that occurred within this time window. Individual cells were isolated, placed into single wells, and incubated in growth conditions. The entire contents of individual wells (30 wells tested per time point) were plated using the plaque assay every half hour, to quantify cell lysis and virus production, starting at 4 hrs and ending at 12 hrs after initial infection (Figure 2A; full details in the Materials and Methods).

**Figure 2:**
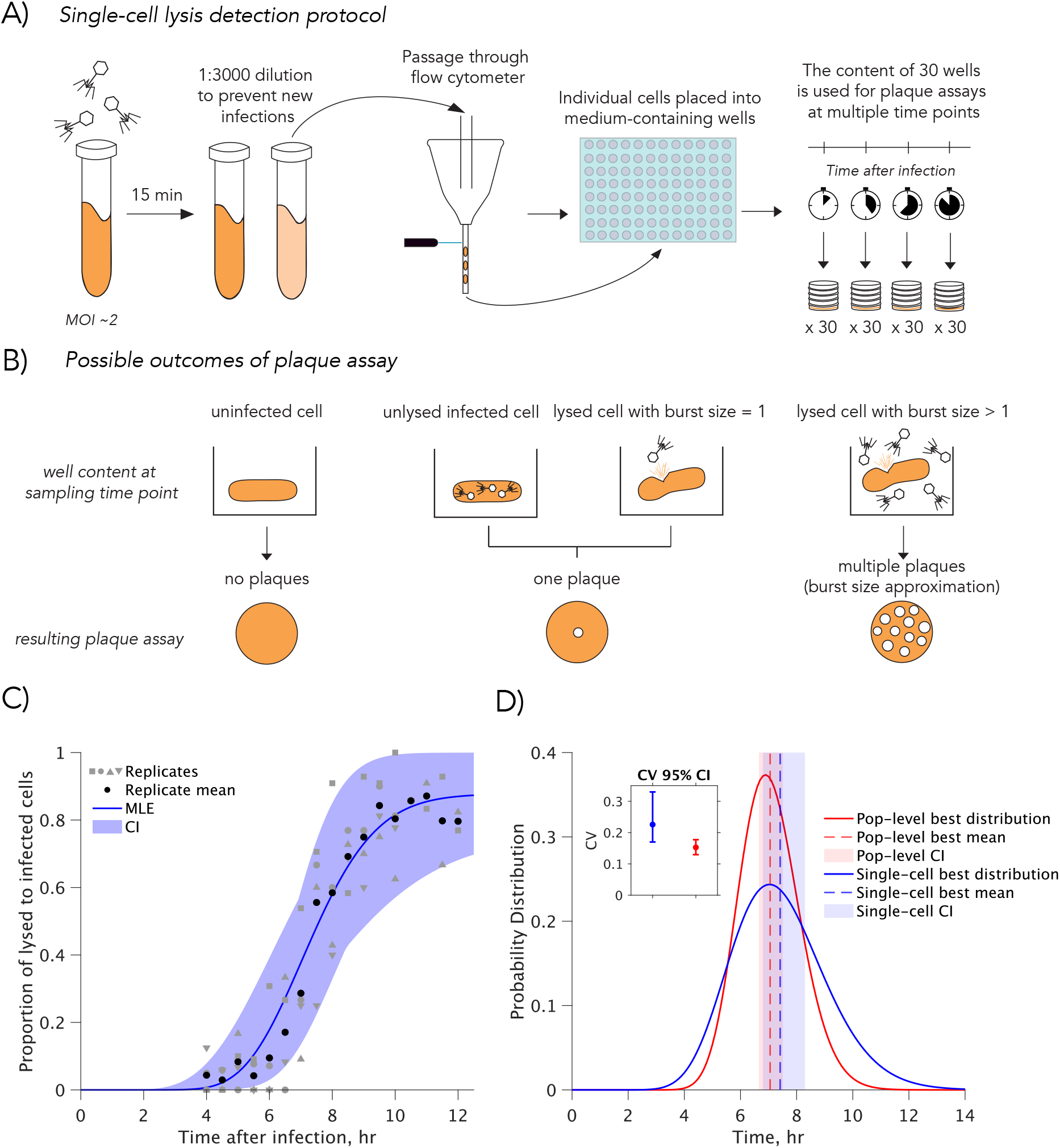
Single-cell heterogeneity in lysis time. **A)** A ‘single-cell lysis detection protocol’ was developed to reconstruct the Cumulative Distribution Function (CDF) of a virus-microbe latent period. A microbial population is inoculated with phage, and after 15 minutes, the population is diluted to prevent further infections. The population is then passed through a flow cytometer, where individual cells are isolated and placed into wells. At multiple time points, the contents of 30 wells are harvested for plaque assays to assess infection status. **B)** At each time point, plaque assays can yield three outcomes: i) no plaques for uninfected cells, ii) a single plaque for cells that were infected but not yet lysed, or for lysed cells with a burst size of 1, and iii) multiple plaques for cells that lysed and released multiple infective viruses. The number of assays with one or more plaques reflects the number of infected cells. Assuming few infections with a burst size of one, the proportion of lysed cells is approximated by the ratio of assays with multiple plaques to the total number of infected cells. **C)** Prediction of the latent period CDF based on 4 experimental replicates of the ‘single-cell lysis detection’ method for all time points from 4 to 10 h, and 2-3 replicates for 10.5 to 12 h (gray symbols). The solid line indicates the best fit, while the shaded region represents the 95% CI. The different-shaped gray symbols indicate the different biological replicates. **D)** The Probability Distribution Function (PDF) of the latent period derived from population-level viral density time series (Figure 1, red) and the Cumulative Distribution Function (CDF) from the single-cell lysis detection method (blue). The CI for both the mean latent period (shaded region) and the coefficient of variation (CV, figure inset) overlap. Table S3 lists all predicted parameter values.

By the time of plating, three scenarios could have occurred within the individual well (Figure 2B): (i) The cell in the well was not productively infected. Under this scenario, the plaque assay would result in no visible plaques. (ii) The cell was infected but did not lyse before the well contents were plated. Here, cell lysis would occur on the plate and all virions would be released from the same location, resulting in a single plaque. (iii) The cell lysed before the well contents were plated. At the time of plating the contents of the well were primarily free virions. When the contents were used for plaque assays, each individual virion would be plated in a different spot, resulting in as many plaques as infective viral particles produced - a measure of that individual cell’s burst size. If the infected cell released a single infective particle, there would be a single plaque. Note that cells that were succesfully infected but not lysed by the plating time point and cells lysed with a burst size of one infective particle would both result in a plaque count of 1. These two scenarios are indistinguishable from each other [34] (Figure 2B).

If we assume that no infections (or a negligible number) result in a burst size of 1, we can approximate the number of cells lysed by the sample time point as the number of plates with 2 or more plaques (scenario iii). We can calculate the total number of successful infections by counting the plates that have 1 or more plaques (adding scenarios ii and iii). Then, the probability that an infected cell had lysed by the sample time point is the ratio between the number of lysed cells (scenario iii) and the total number of infections (the sum of scenarios ii and iii). Following the increase of probability of cell lysis across time gives us the cumulative distribution function (CDF) of the lysis time distribution (Figure 2C).

Using this method, we observed plaque assays with 2 plaques or more at 4 hr after initial infection. The proportion of lysed to infected cells continued to increase until 10 hrs after initial exposure, suggesting that this is the longest lysis time for infected cells within the population. Interestingly, even after 12 hrs, the proportion of lysed to infected cells did not reach 1, raising the possibility that some infections resulted in a burst size as small as 1 infectious viral particle per infected cell (equivalent to one plaque-forming unit) (Figure 2C), as observed previously in the original implementation of the burst-size protocol [34]. By considering burst sizes of 1 (see Materials and Methods, Figure S3), we find that the single-cell latent period data is best described by a Gamma distribution with mean 7.4 hr (CI: [6.8, 8.3]) and CV 0.23 (CI: [0.17, 0.33]) (Figure 2C,D, see Figure S4-6 for evidence on our ability to infer latent period distributions using data from the ‘single-cell lysis detection protocol’). This estimation of latent period heterogeneity is compatible with the range observed in other phage-bacteria (ranging from 0.05 to 0.21 in specific coliphages) [23, 26, 27], a matter we revisit in the discussion.

Critically, when we compare the latent period distribution predicted using our non-linear model based on the population-level time series data to the single-cell experimental data, we find a high degree of congruence. In particular, we find that the predicted average and CV fall within the confidence intervals obtained from the single-cell data (Figure 2D). Similarly, the average burst size is successfully predicted by our model from population-level multi-cycle growth curves (Figure S7). These results show that single-cell heterogeneity of viral life history traits can be predicted from population-level dynamics.

### 2.3 Lysis time variability influences burst size

We next sought to determine whether lysis time is related to burst size at the single-cell level. In addition to characterizing the latent period distribution, the single-cell experimental assay provides a measure of the burst size of each infected cell. At a given plating time *t*, we observe burst sizes from all cells that have lysed before that time. As a result, the experiment captures cumulative rather than instantaneous lysis events, yielding an effective burst size – defined as the average burst size of all cells that have lysed by time *t*. The single-cell experiment shows that the effective burst size increases with sampling time (Figure 3A) and begins to plateau around 10 hours after infection, which coincides with a time after which few new lysis events occur (Figure 2D). These observations suggest that the burst size of individual cells increases with latent period.

**Figure 3:**
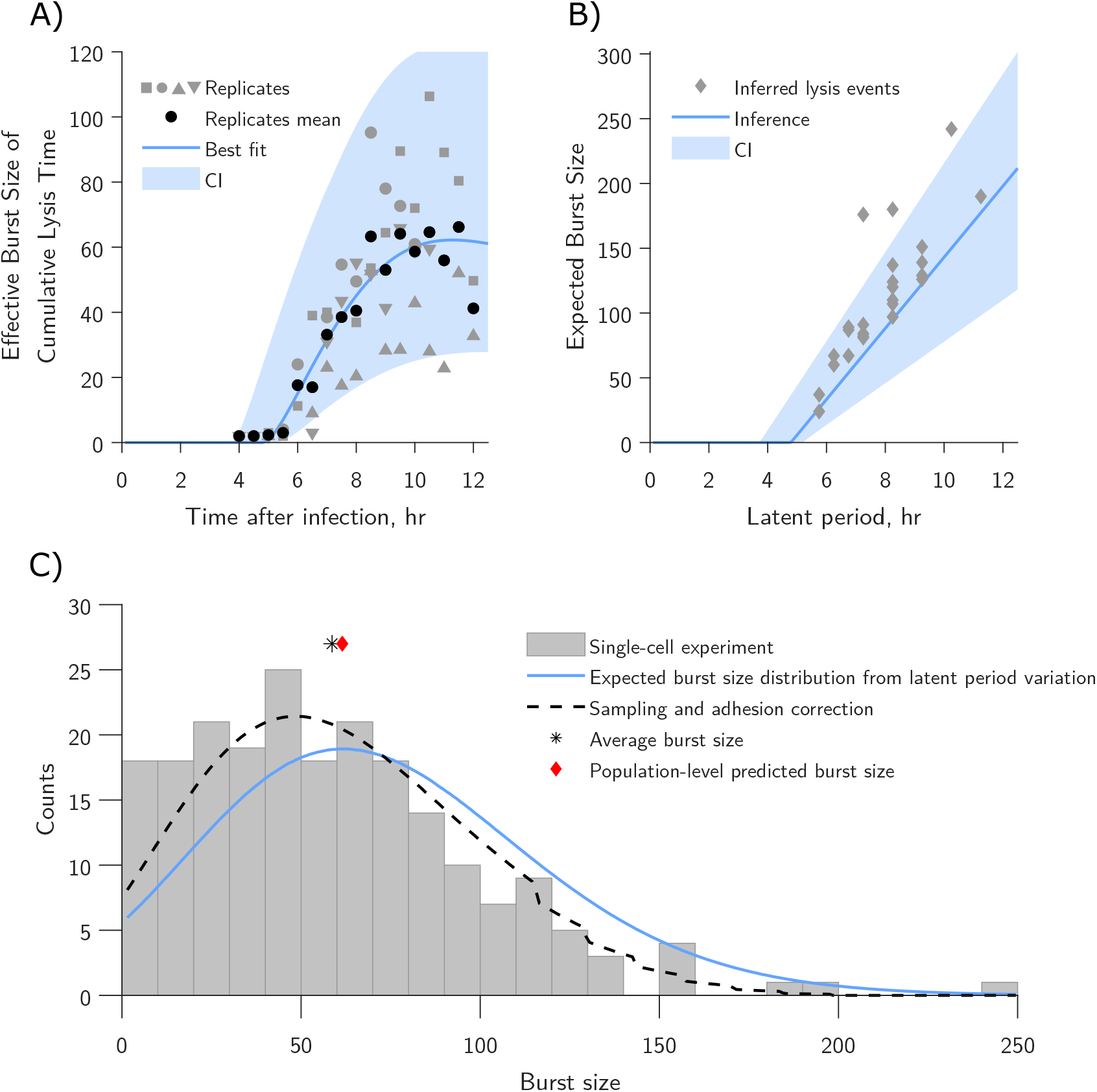
Relationship between burst size and latent period. **A)** The average burst size at each sample time point from the ‘single-cell lysis detection method’ reflects the interplay between the latent period distribution and the burst size-latent period relationship. This relationship is inferred using the previously predicted latent period distribution. The different-shaped gray symbols indicate the different biological replicates. **B)** The predicted relationship shows a linear increase in burst size as a function of latent period, with a rate of progeny production of 27.4 viral particles per hour [CI: 15.0, 34.9], beginning at 4.8 hr [CI: 3.7, 5.2] post-infection. Note that the burst sizes from inferred lysis events (gray diamonds, see Methods) were not used for inference, but support this linear relationship. **C)** The burst size distribution using all plaque assays starting at 9 hr after infection (n = 213, gray bars) is compared to the expected burst size distribution derived from our predictions of latent period PDF and latent period to burst size relationship (solid orange line), and to the distribution we expect to observe when we correct for sampling at different timepoints, and for time-dependent viral particle adhesion to well surfaces (dashed line). The black asterisk represents the average burst size observed from single cell data, the red diamond represents the burst size predicted from the population-level inference (Figure 1). Table S4 lists all predicted parameter values.

The effective burst size at each sampled time point *t* represents a weighted sum of the underlying latent period–burst size relationship, with weights proportional to the lysis probability up to time *t* (see Methods). Using the effective burst size time series, we analyze three models that could potentially describe the relationship between the latent period, *τ* and burst size, *β*(*τ*): (i) a piecewise linear model, (ii) a saturating model represented by a Hill function, and (iii) a logistic growth model that characterizes the lysis time–burst size relationship in chemically induced lysis of individual *λ* lysogens [36]. We find that a piecewise linear model for *β*(*τ*) that incorporates the latent period distribution (Figure 2D) and corrects for time-dependent adhesion of virions to the well surface in our experimental protocol (Figure S8,Materials and Methods), minimizes the mean squared error for the effective burst size across four replicates (Figure 3A). The model predicts that the earliest cells begin to burst around 4.8 hr after infection [CI: 3.7–5.2] with a rate of progeny production of 27.4 infective viral particles per hour subsequently [CI: 15.0, 34.9]. We note that this initial lysis time is consistent with the one-step growth curve inference of 4 to 4.5 hr – again reinforcing the interpretation that conventional latent period assays provide information on the earliest lytic events only.

To further validate predictions of a quantitative relationship between burst size and latent period at the scale of individual cells, we infer which specific lysis events occurred at time intervals of half an hour, given differences in the effective burst size distribution at different sampling times (see Materials and Methods). Our predictions support a linear latent period to burst size model after a delay to first burst (Figure 3B, Figure S9). Note that this linear relationship may only hold in the lysis time interval between 5 to 10 hrs, after which the probability of new lysis events drops significantly. As observed by Kannoly *et al*. [36], long lysis times would result in the depletion of bacterial resources and a latent period to burst size function that flattens at longer times. We note that our model presumes that variability in burst sizes are due exclusively to variability in the post-eclipse period, the time during infection after the first infectious viral particles are formed. Recent work has suggested that variability in the initial stages of infection, i.e. from adsorption to start of capsid production, can account for a substantial proportion – around 89% – of total lysis time variability (re-analysis of data from [23] is shown in Table S5). We extended the present inference framework to include variation in the eclipse period and, again, find that the piecewise linear relationship between burst size and latent period remains the best-supported model (Figure S10, Table S6). The optimal parameter set for this model attributes a relatively, smaller fraction (0.05) of latent period variability to eclipse period variability (Figure S10). The model with eclipse period variability also leads to an improved prediction of the linear burst size-lysis time relationship (see Figure S10D). We caution that the current single-cell protocol does not enable direct observation of the eclipse period and, as a result, the eclipse period is not robustly identifiable (Figure S11). While further work is required to enable robust decomposition of lysis time variability, our results provide the first empirical evidence that heterogeneity in progeny production is a direct result of variability in lysis time (Figure 3C) –shedding light on the source of burst size variability described by Delbrück 80 years ago.

## 3 Discussion

In this study, we characterized the latent period distribution of the cyanophage Syn9 when infecting *Synechococcus* sp. WH8109 using two complementary approaches. First, we applied a population-level modeling framework [20] to infer life-history traits from viral density time series data in multi-cycle growth curve experiments. Second, to measure variability in lysis time, we developed a single-cell lysis detection method that we adapted from previous work [34] to examine single-cell variability in viral progeny production. In addition, this method allowed us to explore the relationship between burst size and latent period at the single-cell level. The single-cell method confirmed that lysis time heterogeneity at the single-cell level can be captured by population-level data. We found that burst size variability is shaped by intrinsic lysis time variability with burst sizes continuing to rise linearly as a function of the latent period – and not saturating within physiologically relevant timeframes.

A wide range of viral systems exhibit intrinsic phenotypic heterogeneity within a population [37–40]. However, quantifying such variability requires performing measurements at the scale of single cells. These limitations have led to the development of population-level methods for characterizing viral life-history traits, e.g., the one-step growth curve [41]. Despite recognition of the presence of heterogeneity, in practice the latent period is often directly inferred from population scale data without accounting for heterogeneity and its impact on dynamics. Here, we leveraged a recently proposed inference method [20] to estimate life-history traits from population-level time series, showing that it is possible to reliably estimate single-cell latent period distributions without using a single-cell assay. This finding makes it possible to link population-level measurements with underlying cellular processes, to reduce biases in conventional latent period estimates, and to facilitate the development of dynamical models that incorporate empirically grounded trait heterogeneity.

The inference of latent period variability yielded an estimated Coefficient of Variation (CV) of 0.23 associated with the environmentally relevant cyanophage Syn9. This finding implies that individual lysis events will most likely occur between 4 and 11 hours post-infection. This variability is compatible with the range previously observed for coliphage systems including *λ* and T7, whose CVs range from 0.05 to 0.12 for *λ* lysogens [26, 27] to 0.21 for lytic T7 infections [23]. The observed variability in lysis time for both coliphage and Syn9 is low compared to other viral processes and traits, such as burst size, DNA injection time, and lysogenization decisions [23, 25, 34, 42]. We also caution that the presence of additional variability may pose problems for population-level inference of single-cell variability. These observations suggest that relatively tight regulation of lysis timing may be a broadly conserved feature across diverse phage-host systems. The molecular mechanisms governing the latent period have typically been characterized in dsDNA phage with two component holin-endolysin lysis systems infecting gram-negative bacteria like *E. coli* [43, 44]. In those systems, holins and anti-holins act as a clock potentially tightly regulating lysis timing [45–48]. In contrast, single-component lysis systems often exhibit greater variability in lysis timing with CV values up to 0.48 [28, 44]. Further work would help elucidate whether a holin-endolysin-based mechanism helps explain the relative low levels of lysis time variability for Syn9. In this study we were able to infer both latent period and burst size variation at single-cell scales. The variability in burst size of infected cells has been recognized since foundational studies of phage biology [19, 25, 34], even as the origins of burst size variability have remained elusive. Burst size variation is likely influenced by factors such as host cell physiology and size, as well as virus-host compatibility, and the virus latent period [19, 23, 49–51]. Previous studies have hypothesized that burst size is positively, linearly related to latent period [52] – a feature commonly incorporated in the study of life-history trait evolution [53–58]. This relationship has support predominantly from population level observations, where phage strains with longer latent periods have larger burst sizes [52]. Here, we provided single-cell evidence that burst size increases linearly with latent period, without signs of saturation, among cells within a population exposed to the same conditions, resulting in variation in burst size being directly linked to variation in latent period. In contrast, our initial population-based inference of the latent period distribution assumed a constant burst size while still providing a good fit to the population-scale data. These findings imply that direct inference of a latent period–dependent burst size relationship may require single-cell burst and timing measurements, or, potentially, an extension of the current population-based modeling framework to account for the age of infection.

In contrast to the present findings of a linear relationship between burst size and latent periods, a recent single-cell study of lysis timing variability within phage *λ* suggested that the rate of progeny production is initially exponential, followed by a plateau as cellular resources are depleted, and did not identify an association between variability in latent period and burst size [36]. In this prior study, lysis was prevented until artificially induced, causing longer latent periods than those that would occur naturally, which led to the emergence of an apparent plateau. Prevention of lysis may have obscured the covariability between lysis time and burst size. Moving forward, it will be important to adapt the current method to understand the relationship between ecological context and trait variability, including in periods of resource limitation in which plateaus may be measurable and ecologically relevant.

Recent work in phage T7 suggests that a significant portion of the variability of the latent period is attributable to variability in the early stages of infection, i.e. before viral capsid production begins [23]. In contrast, in this study, we identified a potentially modest contribution of eclipse period variability to latent period variability (on the order of 5%; Figures S10–S11), albeit robust estimation of the relative magnitude of eclipse period variability is not feasible using our population and single-cell assay protocols. We anticipate that experiments that directly measure the eclipse period in single-cell cohorts will help disentangle the contributions of different processes and clarify patterns of covariation among phage life-history traits in diverse phage-host systems.

The present study finds significant, intrinsic trait variability for infected cells exposed to the same experimental conditions. In natural environments, cyanobacteria are subjected to changing conditions that impact their growth and the expression of viral traits. These include changes in temperature and light availability, including levels and duration over day-night cycles, which impact viral adsorption and production [59–63]. Additionally, resource availability and interactions with other bacterial species and phages may further impact traits (e.g., due to changes in the availability of nitrogen and phosphorus [17, 18]). The inclusion of resource limitation and additional environmental complexity is likely to lead to even greater variability than that observed under a single set of conditions as used in this study. We anticipate that incorporating other forms of ecological feedback into the ‘multi-cycle growth curve’ inference framework will enable estimates of single-cell latent period variability in ecologically relevant conditions. Furthermore, given the generality of the framework, we are optimistic that the underlying mathematical model and single-cell assay can be adapted to other phage-bacteria systems. In particular, the single-cell assay does not rely on the microscope tracking of individual cells leveraged by previous studies characterizing latent period variability [23, 26]. Nonetheless, certain limitations remain. Using a flow cytometer to separate cells can be restrictive, and latent period variability may be difficult to resolve for phage with very short latent periods, since relevant differences may occur on the scale of minutes – an issue that could be addressed via incorporation of microfluidic-enabled plating.

The ecological implications of latent period variability are wide-ranging, influencing both short-term phage–microbe dynamics and long-term evolutionary outcomes. Because variation in the latent period directly affects phage–microbe dynamics [20, 22], it can have significant consequences for viral fitness. Latent period variability can represent both an advantage and a disadvantage for the phage, with earlier lysis events enabling faster reproduction but yielding smaller burst sizes. Since latent period variability appears to be at least partly heritable [26, 27] and produces clear phenotypic effects, latent period variability may represent a trait under selection [24]. For instance, heterogeneity in latent periods may function as a bet-hedging strategy, increasing the likelihood that some infections succeed under fluctuating conditions. Variability can also promote coexistence between phages and their bacterial hosts [64]. More broadly, variation in the latent period shapes the selective landscape of other life-history traits, shifting the average optimal latent period in a population [65] and potentially influencing the evolution of additional traits. Further research is needed to explore whether and how latent period variability stems from a trade-off between burst size and lysis time, either as an unavoidable consequence of molecular stochasticity or as an adaptation strategy [23, 24, 27, 66].

In closing, we have inferred and quantified ecologically meaningful latent period variability in a globally relevant cyanophage. Furthermore, we have shown that it is possible to accurately infer heterogeneity in lysis-associated life history traits at single-cell scales from population-scale dynamics. This inference approach helped determine that variation in the latent period is a major contributor to realized variation in burst size. We anticipate that quantifying the variability of life-history traits in phages will help identify mechanistic principles underlying lysis timing variation and enhance the development of predictive models of viral impacts in therapeutic and environmental contexts.

## 4 Materials and Methods

### 4.1 Bacterial culture growth and phage propagation

*Synechococcus* WH8109 cultures were grown in artificial sea water (ASW) medium [67] with modifications as described in Lindell *et al*. [68], at 21°C and a light intensity of 45 *µ*mol photons m^−2^s^−1^ under a 14:10 light-dark cycle with gentle shaking. The growth rate under these conditions was approximately a doubling a day. Cell density was enumerated using the Influx flow cytometer (BD Biosciences). Cultures of *Synechococcus* were excited with a 457-nm and 488-nm laser and detection based on their orange fluorescence (emission at 580/30 nm) and forward scatter. Yellow-green 1-*µ*m-diameter microspheres (Fluoresbrite) were added to each sample as an internal standard for size and fluorescence. Cell density and culture growth prior to experiments was approximated using chlorophyll *a* fluorescence (excitation at 440 nm, emission at 660 nm) measured in 96-well plates using a Synergy Mx Microplate Reader (Biotek). Phages were propagated by infecting *Synechococcus* WH8109 cultures at a multiplicity of infection (MOI) of ∼0.1. After 24 hours, the lysate was centrifuged at 5467 g for 5 minutes to remove residual host cells and filtered through a 0.22 *µ*m syringe filter (Millex-GV, Millipore) to remove cell debris. The concentration of infective phages was enumerated using the plaque assay. Lysates were diluted and pour-plated in plates containing *Synechococcus* WH8109 at sufficient concentrations to produce lawns. Pour-plating was performed as previously described [69] using ASW medium supplemented with 1 mM sodium sulfite [70] and 0.28% low melting point agarose (Invitrogen) [70, 71].

### 4.2 Virus multi-cycle response curve

Multi-cycle virus-host infection dynamics assays were performed by infecting exponentially growing *Synechococcus* WH8109 cultures (∼ 5 ×10^7^ cells/mL) at a MOI of ∼ 0.1. Samples of 0.1 mL from the infected culture were collected, diluted with 0.9 mL of medium, and filtered through a 0.22 *µ*m syringe filter (Millex-GV, Millipore) to remove host cells. The number of infective phages in the filtrate was then determined by the plaque assay. Four independent biological replicates were performed at different times.

### 4.3 Single-cell lysis detection protocol

To recapitulate the latent period distribution of Syn9 infecting *Synechococcus* WH8109 we conducted a single cell infection experiment. *Synechococcus* WH8109 (1.2 ×10^8^ cells/mL) was infected at an MOI of 2 and diluted 3000-fold after 15 minutes. This was done to limit adsorption time and thus infections, allowing us to measure the latent period at a 15-minute time resolution, and to prevent co-sorting of free phages and cells as previously described [34]. Single cells were then sorted into 96-well plates containing medium using an Influx flow cytometer. To reduce oxidative stress for the sorted cells a heterotrophic helper *Alteromonas* sp. EZ55, was added to the medium in the wells as previously described [34, 72]. The number of infective phages produced by each of the 30 single cells was determined every 30 minutes by the plaque assay in 4 biological replicates. The first was from 4 to 10 h, the second from 4 to 10.5 h and the last two replicates were from 4 to 12 hours, such that there are 4 replicates from 4 to 10 h, 3 replicates for 10.5 hours and 2 replicates from 11 to 12 hours. This was done by plating the entire content of a single well on a plate, as previously described [34]. The four independent biological replicates were performed at different times.

To estimate the adsorption of infective phages to plastic during the experiment we tested changes in titer of a Syn9 lysate in a 96-well plate over an 8 hour period. The Syn9 lysate was diluted with ASW containing helper bacteria to 90 plaque forming units per milliliter. The diluted lysate was aliquoted in 96-well plates and the entire contents of 6 of the wells was assayed periodically by plaque assay. We also measured the number of infective phages prior to putting the lysate in the 96-well plate to quantify the loss of phages from small amounts of liquid loss in the wells.

### 4.4 Predicting life-history traits from population-level time series

We use a modified version of a lysis model accounting for lysis time variability from Dominguez-Mirazo *et al*. [20], described here for clarity. The system of nonlinear differential equations includes susceptible cells, *S*, free viruses, *V* , exposed cells, *E*, and actively-infected cells, *I*. Susceptible cells, *S*, have a maximal cellular growth rate *µ* (hr^−1^) and a total cell population carrying capacity *K*(cells/ml) where 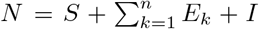 gives the total cell population. We assume that infected cells at any stage of infection do not grow and that cell death rates and viral decay rates are negligible compared to other key rate constants of the system. We assume that viruses and hosts are well-mixed. The rate at which susceptible cells (*S*) are infected is given by: *i*(*t*) = *ϕ S V* , where *ϕ*(ml/hr) denotes the adsorption rate. Infected cells at any stage can adsorb phage at the same rate as susceptible cells without consequences to the ongoing lytic cycle.

We incorporate variability in latent period by assuming that before entering the actively-infected stage, *I*, infected cells advance through several exposed *E* stages: *E*_1_, … , *E*_*n*_, where *n* is a non-negative integer. There are *n* + 1 transitions, and exposed cells (*E*) transition between compartments at a rate of (*n* + 1) *η* with exponentially distributed times. The average time from adsorption (*i*.*e*., entering the first Exposed class, *E*_1_) to cell burst (*i*.*e*., exiting the actively-infected class, *I*) is the latent period mean and equals the inverse of the mean lysis rate, *T* = 1*/η*. At the end of the actively-infected stage (*I*), the cell bursts and free virus (*V*) increases from viral release of *β* virions. The system of nonlinear, ordinary differential equations can be written in the form:

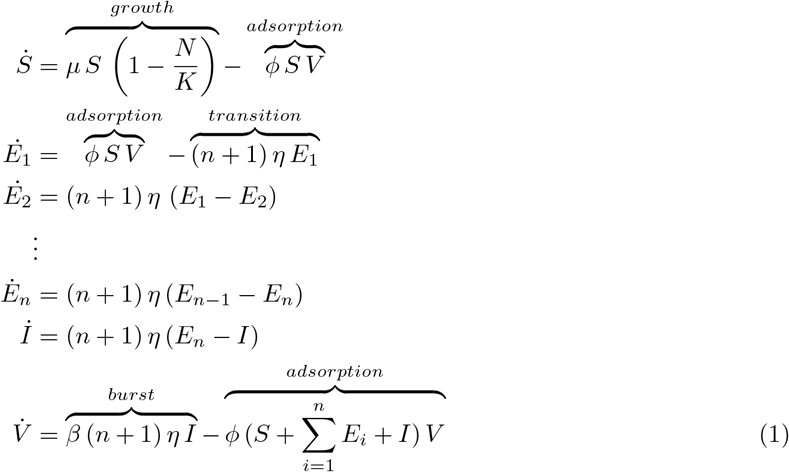

This model describes the latent period distribution as an Erlang distribution with shape *n* + 1 – the number of exposed (*E*) compartments plus the infected (*I*) compartment –, and rate *η* – the lysis rate. In this form, the mean (*T*), variance (*σ*^2^), and coefficient of variation (*σ/T*) of the latent period (LP) are given by:

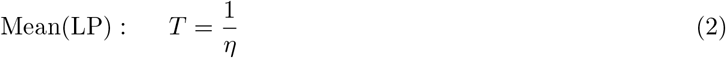

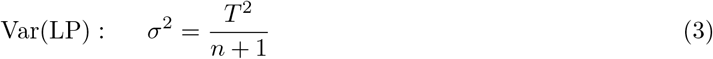

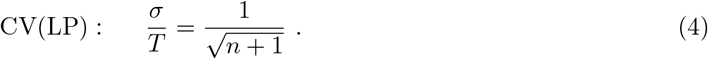

Therefore, the number of *E* compartments dictates the spread of the distribution through the coefficient of variation (CV), with larger *n* leading to tighter distributions and smaller CV values (Figure S1). Since *n* must be an integer, this imposes constraints on the CV values that can be simulated. For example, *n* = 0 corresponds to a CV of 1, and *n* = 1 results in a CV of approximately 0.7. As a result, the model cannot represent CV values between 0.7 and 1. However, latent period distributions with CV below 0.5 can be accurately simulated with a tolerance of 0.05. Based on estimates of lysis timing variability from various studies [23, 26, 28], we anticipate that CV values for latent periods in natural systems will be less than 0.5, aligning with the model’s ability to capture variability in latent period timing.

We leverage a computational pipeline previously designed to fit model parameters to data [20] (see Table S1 for parameter estimates). Briefly, we implement a Markov Chain Monte Carlo (MCMC) algorithm using the Turing package in Julia [73] and inform prior distributions using biological knowledge of the phage bacteria system (see Table S2). We obtain 95% confidence intervals by sampling the MCMC posterior distributions. Convergence analysis can be found in Figure S2.

### 4.5 Predicting latent period distributions from single-cell data

Under our experimental setup, plaque assays at each time point can yield one of three possible outcomes:

1. **No plaques visible:** This outcome indicates the isolated cell was not successfully infected, either because it did not encounter a phage during the co-incubation period or because the infection failed. Previous studies have shown that not all phage-bacteria encounters result in successful infections [34].
2. **One plaque is visible:** A single plaque can result from two types of events: 1) the cell was infected but had not yet lysed by the time the sample was plated. In this case, the infected cell is added to the plate and continues intracellular viral production. When the cell bursts, all virions will be released at a single location in the plate, which results in the formation of a single plaque. 2) The cell was successfully infected and had lysed a single virion (burst size of 1) prior to plating. These two scenarios are indistinguishable under our protocol.
3. **Multiple plaques are visible:** This outcome indicates that cells have lysed and released virions at some point prior to plating. The plaque number represents the burst size of the individual lysed cell.

If we assume the burst size is always larger than 1, the number of plates with multiple plaques reflects the number of infected cells that had lysed by time *t* (denoted as *k*(*t*)), where *t* is the sampling time point. The total number of successful infections observed at each time point (*n*(*t*)) is the sum of scenarios 2 (infected but unlysed cells at the time of plating) and 3 (lysed cells). The proportion of successful infections that had lysed by time *t* (*k*(*t*)*/n*(*t*)) represents the cumulative distribution function of the lysis time distribution. If we were to sample at an infinite time point, we would expect all infected cells to have lysed, resulting in a proportion of 1.

We consider two probability distributions with non-negative support: the log-normal and the Gamma distribution. Note that the Erlang distribution (used to describe latent period distributions in the population-level model) is a special case of a Gamma distribution. Both distributions can be described in terms of the mean lysis time (*T*) and the coefficient of variation (*CV*), as

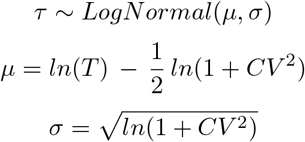

and

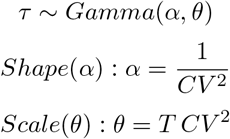

For a combination of latent period mean and CV, we calculate the probability of an infected cell having lysed by the sampling time *p*(*t*), based on the cumulative distribution function (CDF) of either the Gamma or log-normal distribution. The probability of observing our data at each sampling point follows a binomial distribution:

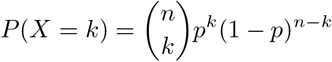

where *k* represents the number of observed lysed cells at the sampling time point, out of *n* infected cells, and *p* is the lysis probability derived from the corresponding latent period CDF.

To account for infections that result in a burst size of 1, we add a parameter *y* where 1™ *y* is the proportion of infected cells we expect to have a burst size of 1. The probability of observing our data at each sampling point follows:

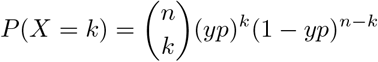

The Maximum Likelihood Estimate (MLE) corresponds to the model (defined by the distribution type (Gamma or log-normal), *y*, mean latent period (*T*), and *CV*) that maximizes the likelihood of observing our data (Figure S3, Table S3). We search for the MLE using the fminsearch MATLAB(R2024a) function, starting the search at multiple combinations of random initial parameters. To calculate parameter confidence intervals, we used profile likelihood methods. In this approach, one parameter is fixed while the confidence interval for the non-fixed parameter is computed. The likelihood ratio is then compared to a chi-square distribution at a one-sided significance level of 0.05. With this experimental protocol and prediction framework, we are able to accurately characterize the latent period distributions for a range of relevant values (Figure S6-7).

### 4.6 Burst size as a function of lysis time

For those plaque assays where the number of plaques is larger than 1, the plaque count reflects the burst size of individual cells. At each sample time point *t*, the plaque count of these assays reflects the observed burst size of cells that lysed by time *t*. The effective burst size at each sampling point that we expect to observe in our protocol is given by,

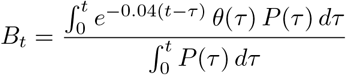

where *P* (*τ*) is the latent period distribution, *θ*(*τ*) is the burst size as a function of the latent period, and the exponential part accounts for time-dependent viral particle adhesion to the well’s surface in our experimental protocol (Figure S8). This equation can be thought of as a sum of expected burst sizes at lysis times shorter than *t*, corrected for particle adhesion, and weighted by the normalized probability of the lysis times.

Leveraging our latent period distribution *P* (*τ*), we evaluate multiple models of expected burst size *θ*(*τ*).

1. Linear model

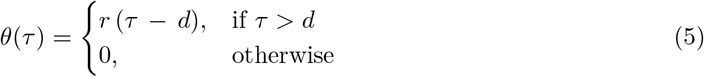

where *r* is the progeny production rate and *d* is the time at which the first cell lyses.
2. Hill function

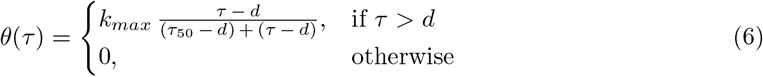

where *d* is the time at which the first cell lyses, *k*_*max*_ is the maximum burst size, and *τ*_50_ is the time at which the expected burst size reaches half of *k*_*max*_.
3. Logistic growth model sourced from [36].

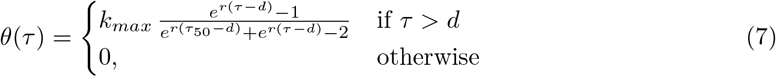

where *d* is the time at which the first cell lyses, *k*_*max*_ is the maximum burst size, and *τ*_50_ is the time at which the expected burst size reaches half of *k*_*max*_.

We find the model that minimizes the Root Squared Mean Error (RSME) using the fminsearch MATLAB(R2024a) function, starting the search at multiple combinations of random initial parameters (Figure S9). We calculate confidence intervals for parameters by bootstraping the data and repeating the search for best parameter combinations, defining a 95% CI as those parameter values that fall within quantiles 0.025 and 0.975.

#### 4.6.1 Inferring lysis events at short time intervals

We use Gaussian Mixture Models to infer the lysis events that occurred in half-hour time intervals. We assume that the probability distribution of burst sizes observed at sample time point *t*_2_, 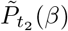, can be described as the collection of burst sizes drawn from two distributions: the burst size distribution of lysis events that occurred by sample point *t*_1_, 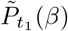, and the burst size distribution of lysis events that occurred in the half-hour interval between time points *t*_1_ and *t*_2_, 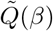, such that

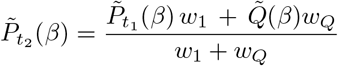

where the weights (*w*) express the probability of lysis events occurring between time 0 and *t*_1_ (*w*_1_), and *t*_1_ and *t*_2_ (*w*_*Q*_) normalized by the probability of lysis events occurring between 0 and *t*_2_ given a latent period distribution *P* (*τ*),

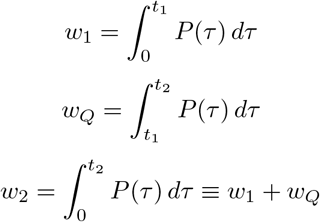

We fit a Gaussian Mixture Model with two components: 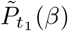 and 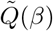. For the first component, we assume a mean and standard deviation equal to the observed burst sizes at time point *t*_1_. For the second component, we initialize the mean and standard deviation estimates to the moments at time point *t*_2_. The initial assumption for the first component and the fit for the second component are used to cluster the burst sizes observed at *t*_2_. The burst sizes that clustered into the second component are lysis events predicted to have occurred in the half-hour interval between *t*_1_ and *t*_2_ as presented in Figure 3B.

## 4.7 Code and data availability statement

We implement the population-level model (Equation 1) and MCMC fitting in Julia v1.7.2 [74] adapted from Dominguez-Mirazo *et al*. [20]. All other analyses were performed in Matlab R2024a [75]. Experimental data and code for simulations and graphics is available at:

https://github.com/WeitzGroup/InferringSingleCellHetereogeneity.git

and archived at:

https://doi.org/10.5281/zenodo.19225909

## 5 Author contributions

MDM, RT, DL and JSW conceived the study. MDM, RT and SK designed the experiments with input from DL and JSW. RT performed experiments with guidance from SK and DL. MDM performed data analysis with guidance from JSW. MDM drafted the manuscript with contributions from all authors.

## 6 Acknowledgments

This research was supported by grants from the Simons Foundation Life Sciences Program 735081 and 529554 to DL and 722153 to JSW. JSW is an investigator at the University of Maryland-Institute for Health Computing, which is supported by funding from Montgomery County, Maryland and The University of Maryland Strategic Partnership: MPowering the State, a formal collaboration between the University of Maryland, College Park and the University of Maryland, Baltimore. The research was supported in part through research cyber infrastructure resources and services provided by the Partnership for an Advanced Computing Environment (PACE) at the Georgia Institute of Technology, Atlanta, Georgia, USA. Funding sources had no role or influence on study design, analysis, interpretation, or submission. We thank Tapan Goel for code review. We thank Hagit Tahan for contributions in creating the illustrations. We thank two anonymous reviewers for comments on the original manuscript that improved this work.

## Notes

https://github.com/WeitzGroup/InferringSingleCellHetereogeneity.git

https://doi.org/10.5281/zenodo.19225909

### Competing Interest Statement

The authors have declared no competing interest.

### Summary of Updates

New supplemental material has been added and corresponding conclusions incorporated into main text.

