## Supplementary Information for "Inferring single-cell heterogeneity of bacteriophage lysis-associated life-history traits from population-scale dynamics"

March 26, 2026

### Supplementary Text

#### S1 Population-level trait inference

To facilitate the reading of the supplementary material, we include the model description found in the main text. We use a modified version of a lysis model accounting for lysis time variability from [1]. A parameter explanation can be found in Table S1.

$$
\begin{aligned}
\dot{S} &= \overbrace{\mu\, S\left(1-\frac{N}{K}\right)}^{growth} - \overbrace{\phi\, S\, V}^{adsorption}\\
\dot{E}_1 &= \overbrace{\phi\, S\, V}^{adsorption} - \overbrace{(n+1)\,\eta\, E_1}^{transition}\\
\dot{E}_2 &= (n+1)\,\eta\,(E_1 - E_2)\\
&\vdots\\
\dot{E}_n &= (n+1)\,\eta\,(E_{n-1} - E_n)\\
\dot{I} &= (n+1)\,\eta\,(E_n - I)\\
\dot{V} &= \overbrace{\beta\,(n+1)\,\eta\, I}^{burst} - \overbrace{\phi\,(S + \sum_{i=1}^{n} E_i + I)\, V}^{adsorption}
\end{aligned}
\tag{1}
$$

This model describes the latent period distribution as an Erlang distribution with shape $n+1$ – the number of exposed ($E$) compartments plus the infected ($I$) compartment –, and rate $\eta$ – the lysis rate. In this form, the number of E compartments determines the spread of the distribution (Figure S1), with the mean ($T$), variance ($\sigma^2$), and coefficient of variation ($\sigma/T$) of the latent period given by:

$$\text{Mean(LP)}: \quad T = \frac{1}{\eta} \tag{2}$$

$$\text{Var(LP)}: \quad \sigma^2 = \frac{T^2}{n+1} \tag{3}$$

$$\text{CV(LP)}: \quad \frac{\sigma}{T} = \frac{1}{\sqrt{n+1}}\,. \tag{4}$$

##### S1.1 Details on parameter inference

In this section we describe in more detail the parameter inference found in the methods of the main text and adapted from [1]. Using population-level timeseries we fit data of four experimental replicates to our compartmental model of lytic infections to find the parameter values that allow the model to best describe the experimental data. Host-only parameters (growth rate, $\mu$, and carrying capacity, $K$) are inferred by fitting the model to bacterial count time series performed in the absence of virus, simulating a null viral initial density. In the next step, $\mu$, and $K$ are fixed to the inferred values and the host-phage pair traits ($\phi$, adsorption rate; $\beta$, burst size; $T$, latent period mean; $CV$, latent period coefficient of variation) are estimated, along with bacterial and viral initial densities ($S_0$ and $V_0$), using viral density time series. A parameter description and estimated values are found in Table S1.

Inference is done using Markov Chain Monte Carlo (MCMC) implemented in the probabilistic inference package Turing [2] in the Julia language v1.8 [3]. MCMC is a class of algorithms that aim to obtain the equilibrium probability distribution of the model parameters [4]. The prior distributions used in the MCMC are modeled as a LogNormal distribution, $LogNormal(\theta,\nu)$, where $\theta$ and $\nu$ are defined as follows: $LogNormal(\theta,\nu)$, where $\theta = \ln mode + \frac{2}{3}\ln\frac{mean}{mode}$ and $\nu = \sqrt{\frac{2}{3}\ln\frac{mean}{mode}}$. Here, the mode is a point estimate derived from biological expectations, while the mean is set to twice the mode value (Table S2).

We use the No-U-Turn Sampler (NUTS) from the Turing package [2] to sample the posterior distribution. For each implementation, we ran MCMC chains for 2000 iterations with a 1000-iteration warm-up period. The target acceptance ratio was set to 0.45. Since chains may converge to different parameter ranges, we select the set of chains that converge to consistent values across all parameters and minimize

| Parameter | Point estimate | 95% CI | Unit |
|---|---|---|---|
| $\mu$, growth rate | 0.03 | [0.02, 0.06] | $hr^{-1}$ |
| $K$, carrying capacity | $3 \times 10^8$ | $[1 \times 10^8, 7.7 \times 10^8]$ | CFU/ml |
| $\phi$, adsorption rate | $1.6 \times 10^{-8}$ | $[1.3 \times 10^{-8}, 2 \times 10^{-8}]$ | ml/(CFU×hr) |
| $\beta$, burst size | 61.5 | [49, 76] | |
| $\eta$, lysis rate | 0.14 | [0.13, 0.15] | $hr^{-1}$ |
| CV, coefficient of variation | 0.15 | [0.13, 0.18] | |
| n, number of $E$ compartments | 43 | [30, 58] | |
| Initial condition | Point estimate | 95% CI | Unit |
| $S_0$, initial microbial density | $4.1 \times 10^7$ | $[3.6 \times 10^7, 4.7 \times 10^7]$ | CFU/ml |
| $V_0$, initial free virus density | $3.9 \times 10^6$ | $[3.1 \times 10^6, 5 \times 10^6]$ | PFU/ml |

Table S1: **Population-level parameter estimates.** The parameters and initial conditions correspond to those in the dynamical model of lytic infections. The corresponding fits to the data are shown in main Figure 1.

| Parameter | Parameter description | Units | Prior distribution | Prior mode | Prior bounds |
|---|---|---|---|---|---|
| $\mu$ | Growth rate | $CFU/ml$ | $LogNormal(\theta, \nu)$ | 0..03 | [0, 2] |
| K | Carrying capacity | $CFU/ml$ | $LogNormal(\theta, \nu)$ | $9 \times 10^7$ | $[10^3, 10^{10}]$ |
| $\phi$ | Adsorption rate | $ml/(CFU \times hr)$ | $LogNormal(\theta, \nu)$ | $5 \times 10^{-9}$ | $[10^{-10}, 10^{-7}]$ |
| $\beta$ | Burst size | | $LogNormal(\theta, \nu)$ | 40 | [5, 100] |
| $\eta$ | Lysis rate | $hr^{-1}$ | $LogNormal(\theta, \nu)$ | 0.14 | [0.083, 0.3] |
| CV | Coefficient of variation | | $LogNormal(\theta, \nu)$ | 0.2 | [0.03, 1] |
| $\sigma_{loglikelihood,h}$ | Error chain for host series | | $InvGamma(1, 0.5)$ | NA | [0, 2] |
| $\sigma_{loglikelihood,v}$ | Error chain for viral series | | $InvGamma(1, 0.5)$ | NA | [0, 2] |

Table S2: **MCMC priors used for parameter inference.** The specific values for $\theta$ and $\nu$ are described in the text above.

the error chain. To prevent high autocorrelation, we achieved chain convergence by thinning chains to obtain 3000 steps. We obtain 95% confidence intervals by sampling the resulting posterior distributions.

To verify convergence of the MCMC chains, we computed the potential scale reduction factor, $\hat{R}$ [5], for the pooled chains. The $\hat{R}$ value for each parameter quantifies the ratio between the average variance across different chain samples and the total variance of the entire chain. When the chains have successfully converged to a posterior distribution, $\hat{R}$ will be close to 1. Conversely, if convergence has not occurred, $\hat{R}$ values will exceed 1. Our calculated $\hat{R}$ values for all chains are consistently below 1.1, indicating that convergence has been achieved. We calculated the Effective Sample Size (ESS, $N_{eff}$) ratio by dividing $N_{eff}$ by the total number of iterations in the pooled chain ($N = 3000$). A low ratio, typically defined as $\frac{N_{eff}}{N} << 0.1$, indicates potential sampling problems and unreliability in estimating confidence intervals. The ESS ratio for all of our chains is near 0.1. Both $\hat{R}$ and $N_{eff}$ were computed using the `ess_rhat` function from the `MCMCDiagnosticTools` Julia package [6] (Figure S2).

### S2 Single-cell level trait inference

To enhance clarity in interpreting the predicted model and parameters, we have included the equation from the main text in this supplementary material section. For a combination of latent period mean and CV, we calculate the probability of an infected cell having lysed by the sampling time $p(t)$, based on the cumulative distribution function (CDF) of either a Gamma or log-normal distribution. The probability of observing our data at each sampling point follows a binomial distribution:

$$P(X = k) = \binom{n}{k} (yp)^k (1 - yp)^{n-k}$$

where $k$ represents the number of observed lysed cells at the sampling time point, out of $n$ infected cells, $p$ is the lysis probability derived from the corresponding latent period CDF, and $y$ is the proportion of infected cells we expect to have a burst size larger than 1. The Maximum Likelihood Estimate (MLE) corresponds to the model that maximizes the likelihood of observing our data.

#### S2.1 Selecting a latent period distribution model

We explored four models: a Gamma and a lognormal distribution with parameter $y$ fixed to equal 1 (all burst sizes are expected to be larger than 1), and a Gamma and a lognormal distribution estimating parameter $y$ (Figure S3). As discussed in the main text, the model that maximizes the likelihood is a Gamma distribution with mean 7.4 hr, CV of 0.23, and $y = 0.88$ (Table S3). The estimate of $y$ echoes the one observed previously for the same system for infected cells 24 hr after infection [7].

| Parameter | MLE [95% CI] | | | |
|---|---|---|---|---|
| | Gamma, $y = 1$ | **Gamma, estimated** $y$ | LogNormal, $y = 1$ | LogNormal, estimated $y$ |
| $T$ ($hr^{-1}$), latent period mean | 8.1 [7.7, 8.5] | 7.4 [6.8, 8.3] | 8.1 [7.7, 8.6] | 7.5 [6.8, 8.6] |
| CV, coefficient of variation | 0.29 [0.24, 0.35] | 0.23 [0.17, 0.33] | 0.3 [0.25, 0.38] | 0.25 [0.18, 0.38] |
| $y$, proportion of lysed cells with burst size larger than 1 | | 0.88 [0.75, 1] | | 0.9 [0.76, 1] |

Table S3: **Single-cell parameter estimates.** Parameters estimated for latent period distribution based on single-cell detection protocol experimental data (Figure S3). The model with maximum likelihood is a gamma distribution with estimated y (marked in bold). The corresponding fits to the data are shown in Figure S3, and the best model (marked in bold) is shown in main text Figure 2.

#### S2.2 Assessing prediction accuracy

To show that we can accurately predict latent period distributions from data derived from the single-cell lysis detection protocol, we simulated the experiment and evaluated how well we could predict the underlying latent period distribution used in the simulation. We assumed a probability of 0.4 that a well taken at each time point contains an infected cell, since this is the mean probability observed in Figure S4. We used this probability to sample a binomial distribution and obtain a number of wells that contain an infected cell. Then, for a range of Gamma latent period distributions with latent period mean between the intervals 5 and 9 hrs, and a CV between 0.05 and 0.4, we sampled the latent period distribution to obtain simulated lysis times for the infected cells. We simulated the single-cell lysis detection protocol as performed, i.e. by taking 30 wells every half-hour starting at 4hr and ending at 10 hr after infection, and checking whether the infected cells would have lysed by the time of sampling. We assumed that about 12% of infected cells result in a burst size of 1 (Figure S3, Table S3). Notably, this corresponds well to experimental data that previously reported that 11% of cells infected by Syn9 produced a single plaque after overnight incubation [7]. Using this simulated experiment, we predicted the latent period distribution as explained in the methods section of the main text. We repeated this simulation 100 times and observed our ability to accurately predict the latent period distribution. We consider the distribution to be successfully predicted if the maximum likelihood estimate of the mean latent period lies within 1 hr from the underlying mean with which the experiment was simulated, and if the estimated CV lies within 0.1 of the original one (Figure S5).

We evaluated the impact of incubation time, i.e. the time allowed for phages to adsorb to cells before dilution and sorting in the single-cell lysis detection protocol, on the observed lysis time distributions. To do so, we simulated cell lysis for incubation periods ranging from 5 to 30 minutes, using the adsorption rate from the population-level parameter inference (Table S1) and the latent period distribution parameters from the single-cell estimation (Figure S3). Lysis times were sampled by adding a random infection time with an exponentially distributed adsorption time, and a Gamma-distributed latent period. We then compared the mean and CV of the simulated lysis times to the latent period distribution parameters used to generate the data. Our decision to incubate for 15 minutes has a minimal impact on the observed lysis times, affecting the average lysis time by less than 10 minutes – an insignificant change considering the expected average latent period is around 7 hours. The coefficient of variation (CV) is also minimally affected (Figure S6).

### S3 Latent period to burst size relationship

To simplify the reading of this section we have included the model information found in the Methods section of the main text. The effective burst size at each sampling point that we expect to observe in our protocol is given by,

$$B_t = \frac{\int_0^t e^{-0.04(t-\tau)}\, \theta(\tau)\, P(\tau)\, d\tau}{\int_0^t P(\tau)\, d\tau}$$

where $P(\tau)$ is the latent period distribution, $\theta(\tau)$ is the burst size as a function of the latent period, and the exponential part accounts for time-dependent viral particle adhesion to the well's surface in our experimental protocol (Figure S8).

#### S3.1 Selecting a model of latent period to burst size relationship

We evaluate multiple models of expected burst size $\theta(\tau)$.

1. Linear model

$$\theta(\tau) = \begin{cases} r\,(\tau\, -\, d), & \text{if } \tau > d \\ 0, & \text{otherwise} \end{cases} \tag{5}$$

where $r$ is the progeny production rate and $d$ is the time at which the first cell lyses.

2. Hill function

$$\theta(\tau) = \begin{cases} k_{max}\, \frac{\tau - d}{(\tau_{50} - d) + (\tau - d)}, & \text{if } \tau > d \\ 0, & \text{otherwise} \end{cases} \tag{6}$$

where $d$ is the time at which the first cell lyses, $k_{max}$ is the maximum burst size, and $\tau_{50}$ is the time at which the expected burst size reaches half of $k_{max}$.

3. Logistic growth model sourced from [8].

$$\theta(\tau) = \begin{cases} k_{max}\, \frac{e^{r(\tau-d)}-1}{e^{r(\tau_{50}-d)}+e^{r(\tau-d)}-2} & \text{if } \tau > d \\ 0, & \text{otherwise} \end{cases} \tag{7}$$

where $d$ is the time at which the first cell lyses, $k_{max}$ is the maximum burst size, and $\tau_{50}$ is the time at which the expected burst size reaches half of $k_{max}$.

As discussed in the main text, the best parameters for each model were selected as the ones that minimize the Root Mean Squared Error (RMSE), and are shown in Table S4.

| Parameter | RMSE estimate [95% CI] | | |
|---|---|---|---|
| | **Linear model** | Hill function | Logistic growth [8] |
| $d$ ($hr$), time of first burst | 4.8 [3.7, 5.2] | 5.4 [4.7, 5.7] | 1.5 [1, 6.6] |
| $r$ ($hr^{-1}$), progeny production rate | 27.4 [15, 34.9] | | 2.1 [1.1, 5.5] |
| $K_{max}$, maximum burst size | | 118.9 [69.5, 174.7] | 88.2 [67.1, 110.2] |
| $\tau_{50}$ ($hr$), time at which expected burst size reaches half of $k_{max}$ | | 6.7 [5.8, 8.4] | 6.4 [5.1, 6.9] |
| RMSE | 38.6 | 42.4 | 45.6 |

Table S4: **Burst size as a function of lysis time, parameter estimates.** Parameters estimated for burst size to lysis time relationship based on single-cell detection protocol experimental data. The linear model minimizes the Root Mean Squared Error (marked in bold). The corresponding fits to the data are shown in Figure S9 for all models, and the data fit to the linear model is depicted in main text Figure 3.

#### S3.2 Contribution of early infection events to latent period variability

Variability in early events of infection, before the production of infectious viral particles, can contribute to latent period variability. In a prior study, Wedd et al. [9] tracked individual *E. coli* cells infected with phage T7 and quantified the variability of multiple infection processes. Using digitized data from Figure 6 of Wedd et al. [9], we estimate that variability in the early events of infection, i.e. from adsorption to the start of viral protein production, accounts for approximately 89% of the total lysis time variability in phage T7 (see full details in Table S5).

| Time interval | Time interval description | mean (min) | n | CV | $\sigma^2$ | source in [9] |
|---|---|---|---|---|---|---|
| $t_0$ to $t_5$ | adsorption to lysis (latent period) | 18.8 | 23 | 0.21 | 15.59 | Figure 6b |
| $t_3$ to $t_5$ | protein production start to lysis | 7.7 | 160 | 0.17 | 1.71 | Figure 6b |

Table S5: **Contribution of early infection events to latent period variability in phage T7.** Timing of T7 bacteriophage infection cycle processes at the single-cell level were tracked by Wedd et al. [9] and appear in Figure 6b. We calculated the variance of early events during infection, i.e. from adsorption to protein production start ($t_0$ to $t_3$) using the mean and coefficient of variation (CV) stated in the figure caption. Similarly, the variance of the latent period ($t_0$ to $t_5$) was estimated using the reported mean and CV in the figure caption.

#### S3.3 Incorporating eclipse period into the latent period to burst size relationship

Our experimental protocol for measuring latent period variability in cyanophage Syn9 does not directly account for variability in the eclipse period. Instead, we extended the burst size model shown in main Figure 3 to account for the eclipse period. The latent period can be divided into two stages. The first stage, the eclipse period, is the time from entry of the virus into the cell until the formation of the first infectious virus particles inside the cell. The second stage, the post-eclipse period, is a period of continued virus production and is prior to cell lysis and release of virus particles into the environment. The previous model assumes that variability in the latent period arises entirely from the post-eclipse period. We extend this framework by incorporating an explicit eclipse period into the model, as explained below.

We assume that the latent period of an individual cell is composed of two independent periods: the eclipse period (of duration $\epsilon$), which includes all infection steps before the first infectious viral particles were formed, and the post-eclipse period (of duration $p$), during which infectious viral particles accumulate inside the cell. For cells with eclipse period $\epsilon$, the average burst size, $\bar{\beta}_\epsilon$ contributing at observation time $t$ is:

$$\bar{\beta}_\epsilon(t) = \frac{\int_0^{t-\epsilon} e^{-0.04(t-(\epsilon+p))}\,\theta(p)\,Q(p)\,dp}{\int_0^{t-\epsilon} Q(p)\,dp}. \tag{8}$$

Here, $Q(p)$ denotes the probability distribution function of the post-eclipse period and $\theta(p)$ gives the burst size as a function of the post-eclipse period. The exponential term accounts for time-dependent viral particle adhesion to the surface of the well in the experimental protocol (Figure S8), where the decay depends on the time between lysis and measurement, or $t-(\epsilon+p)$. The effective burst size at each sampling point ($t$) that we expect to observe in our protocol is then given by

$$B(t) = \frac{\int_{t_0}^{t} \bar{\beta}_\epsilon M(\epsilon)\,d\epsilon}{\int_{t_0}^{t} M(\epsilon)\,d\epsilon}, \tag{9}$$

where $M(\epsilon)$ denotes the probability distribution function of the eclipse period and $t_0$ is the minimum viable eclipse period, such that $M(\epsilon) = 0\ \forall \epsilon < t_0$. The equation for $B(t)$ can be thought of as a sum of expected burst sizes at lysis times shorter than $t$, corrected for particle adhesion, and weighted by the normalized probability of the eclipse and post-eclipse periods.

It follows that the expected burst size of latent period $\tau$, $\beta(\tau)$, is given by:

$$\beta(\tau) = \frac{\int_0^{\tau} M(\epsilon)\,\theta(\tau-\epsilon)\,d\epsilon}{\int_0^{\tau} M(\epsilon)\,d\epsilon}, \tag{10}$$

where we assume that the latent period distribution $P(\tau)$ is a convolution of the eclipse and post-eclipse periods where $M(\epsilon)$ and $Q(p)$ are Gamma distributed with:

$$\mu(\tau) = \mu(\epsilon) + \mu(p),$$
$$\sigma^2(\epsilon) = v\,\sigma^2(\tau),$$
$$\sigma^2(p) = (1-v)\,\sigma^2(\tau),$$

and where $v$ is the proportion of the latent period variance explained by variance in the eclipse period. We evaluate multiple models of expected burst size $\theta(p)$.

1. Linear model

$$\theta(p) = r\,p \tag{11}$$

where $r$ is the progeny production rate.

2. Hill function

$$\theta(p) = k_{max}\,\frac{p}{p_{50}\,+\,p} \tag{12}$$

where $k_{max}$ is the maximum burst size, and $p_{50}$ is the time at which the expected burst size reaches half of $k_{max}$.

3. Logistic growth model sourced from [8].

$$\theta(p) = k_{max}\,\frac{e^{r(p)}-1}{e^{r(p_{50})}+e^{r(p)}-2} \tag{13}$$

where $k_{max}$ is the maximum burst size, and $p_{50}$ is the time at which the expected burst size reaches half of $k_{max}$.

As part of model selection, we identified the parameters that allow the model to best describe the data. Specifically, we fit the expected burst size model parameters ($r$, $k_{\text{max}}$, $p_{50}$) as well as $\mu(\epsilon)$ and $v$. The parameter $v$, representing the proportion of latent period variability explained by the eclipse period, was allowed to vary between 0 and 1. As before, we find the model parameters that minimize the Root Mean Squared Error (RMSE). The best parameters for each of the models are shown in Table S6 and fits to the data are shown in Figure S10. The linear model with $v = 0.05$ minimizes the RMSE. Nonetheless, the proportion of latent period variability explained by the eclipse period ($v$) cannot be determined robustly from our data (Figure S11).

| Parameter | RMSE estimate | | |
|---|---|---|---|
| | **Linear model** | Hill function | Logistic growth [8] |
| $v$ $(hr)$, proportion of latent period variability explained by eclipse period variability | 0.05 | 0.05 | 0.95 |
| $\mu(\epsilon)$ $(hr)$, average eclipse period | 5.3 | 6.3 | 3.6 |
| $r$ $(hr^{-1})$, progeny production rate | 36.8 | | 5.1 |
| $K_{max}$, maximum burst size | | 195.3 | 458.3 |
| $\tau_{50}$ $(hr)$, post-eclipse period at which expected burst size reaches half of $k_{max}$ | | 1.1 | 4.4 |

Table S6: **Burst size as a function of post-eclipse period, parameter estimates.** Parameters estimated for burst size to post-eclipse period relationship based on single-cell detection protocol experimental data. The linear model minimizes the Root Mean Squared Error (marked in bold). The corresponding fits to the data are shown in Figure S10B.

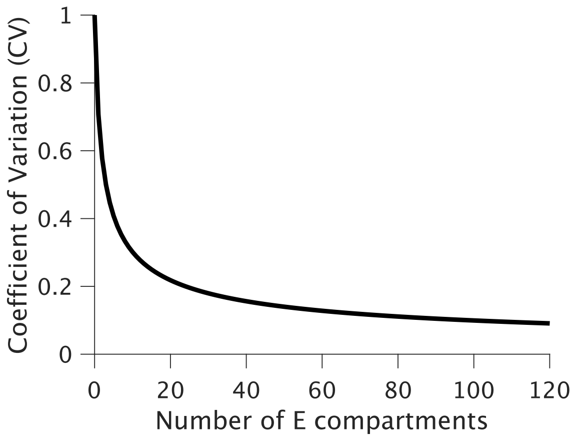

Figure S1: **Relationship between number of E compartments and latent period distribution CV in lytic infection model**. The transition time between intermediate infection compartments is modeled as exponentially distributed. The addition of multiple compartments results in an Erlang distributed latent period distribution. The number of compartments determines the CV of the distribution as $CV = 1/\sqrt{n+1}$. Figure modified from Dominguez-Mirazo *et al.* [1].

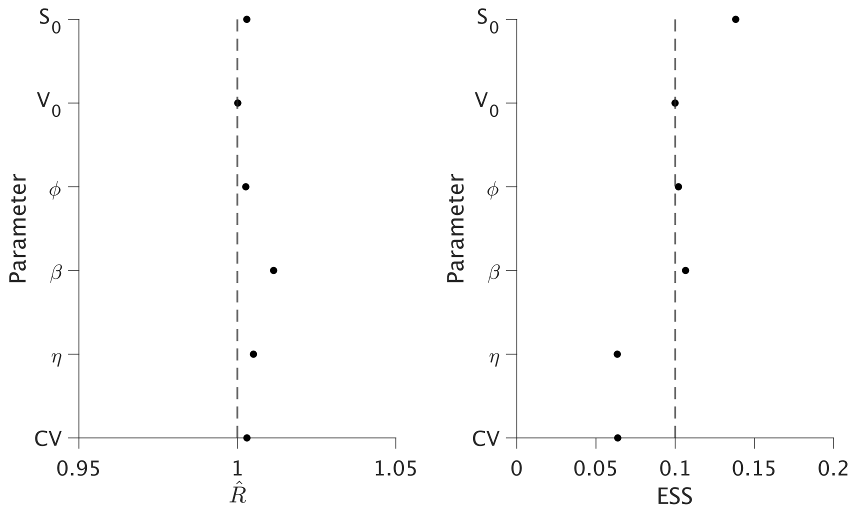

Figure S2: **Chain convergence analysis**. We computed the potential scale reduction, $\hat{R}$ (left panel), and the Effective Sample Size ratio (ESS ratio, right panel) for the MCMC chain. If the chains haven't converged to a single distribution, $\hat{R}$ will be greater than 1. A low ESS ratio, with a common empirical threshold of $\frac{N_{eff}}{N} < 0.1$, indicates potential sampling problems and suggests the estimates of confidence intervals may be unreliable.

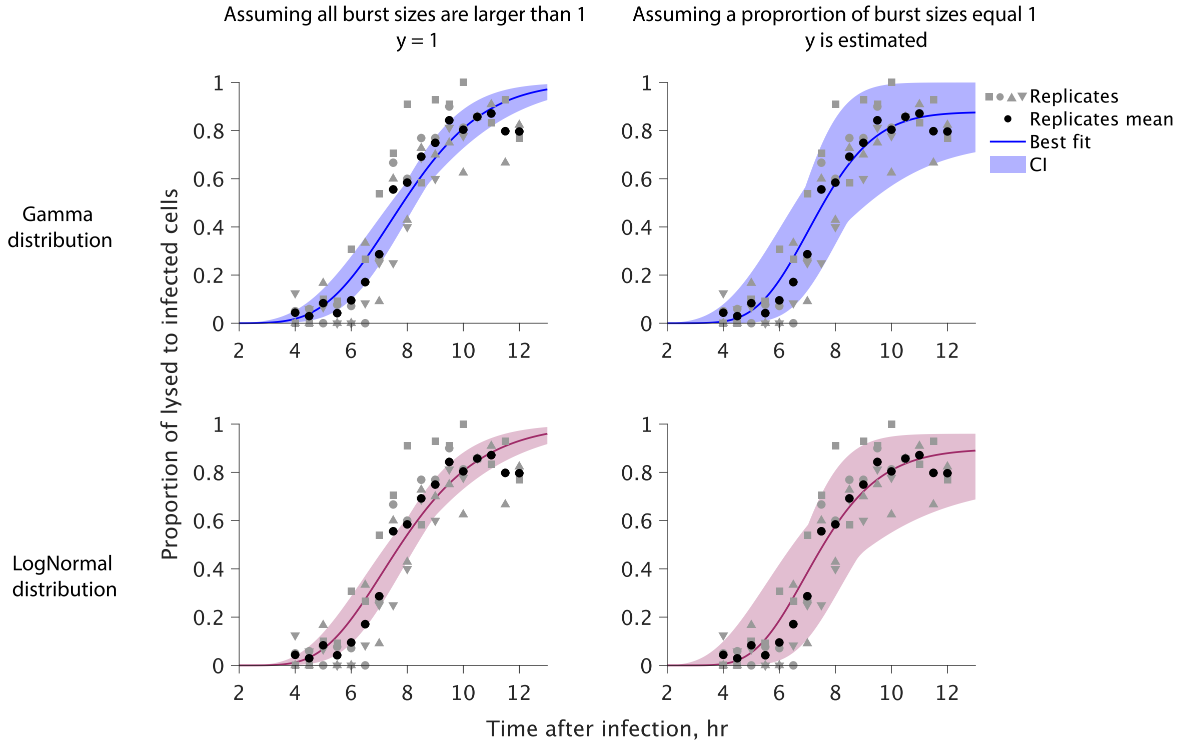

Figure S3: **Selecting best model for CDF estimation from the single-cell lysis detection protocol**. Maximum likelihood data fits for the single-cell lysis detection protocol, assuming a gamma (top row, blue) or log-normal (bottom row, pink) distribution for the latent period. The left column assumes that single-cell burst sizes are always greater than 1 (with parameter $y$ fixed at 1), while the right column allows $y$ to be estimated. The model with the highest likelihood is a gamma-distributed latent period with $y$ smaller than 1 (top right corner, Table S3). This fit is depicted in main text Figure 2.

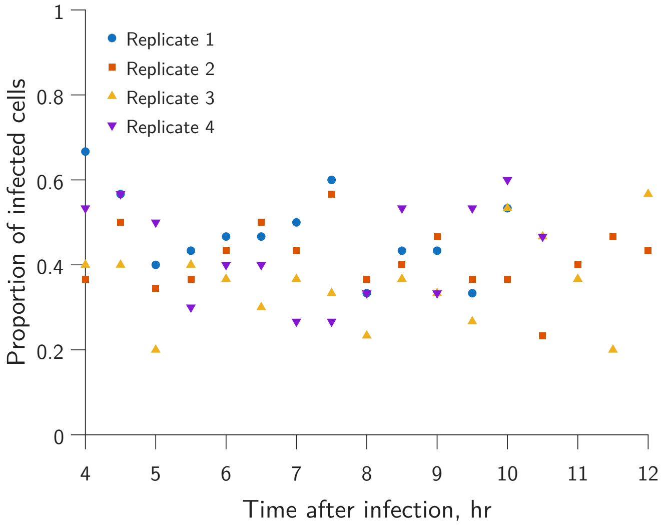

Figure S4: **Sampling time does not affect infection probability**. Proportion of infected cells across time for wells sampled at multiple time points. Cells are assumed to be infected if the sampling contents result in one or more plaques in a plaque assay. For each replicate, the content of approximately 30 wells is used for plaque assays at each time point. The proportion of infected cells is consistent across sampling times.

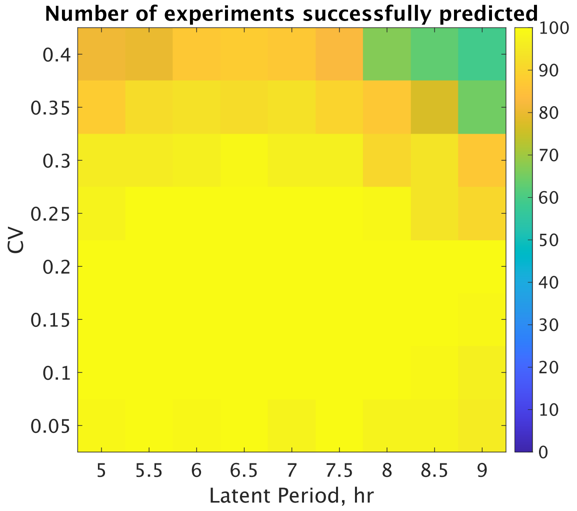

Figure S5: **Accuracy of CDF prediction for relevant range of parameters.** By simulating the single-cell lysis protocol assuming a large range of latent period distributions, we show that the CDF prediction framework we employed can accurately determine latent period distributions from the experimental data. The color in the heatmap represents the number of experiments (out of 100) for which the maximum likelihood estimate of the latent period distribution mean falls within 1 hour, and the CV falls within 0.1, the underlying parameters with which the experiment was simulated.

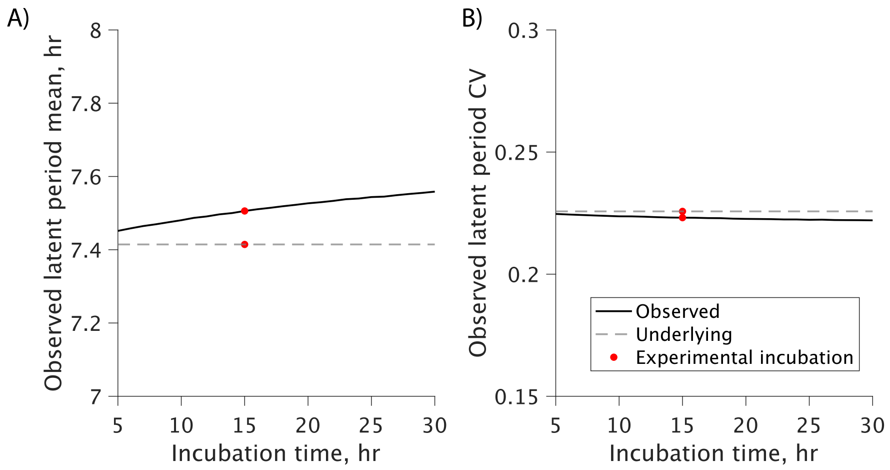

Figure S6: **Effect of incubation time on observed latent period distributions.** We simulate lysis times by randomly sampling infection times, assuming an adsorption rate predicted by the population-level estimation (Table S1) and a gamma-distributed latent period based on the single-cell estimation (Figure S3). Incubation times range from 5 to 30 minutes. As incubation time increases, the average lysis time also increases (A), while the coefficient of variation (CV) decreases (B). The 15-minute incubation time used in the 'single-cell detection protocol', i.e. the time allowed for adsorption prior to dilution and sorting, (red dot) results in an average lysis time that is less than 10 minutes longer than the average latent period used for data generation (dashed line), with minimal impact on the CV.

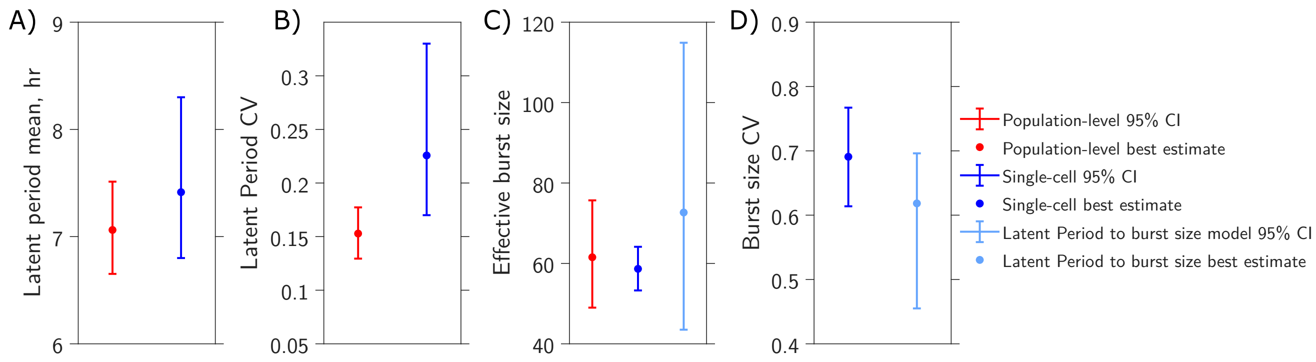

Figure S7: **Comparing predictions for population-level and single-cell level estimates.** We compared estimates of the latent period mean (A), latent period Coefficient of Variation (CV) (B), effective burst size (C), and burst size CV (D) for the population-level (red, main text Figure 1), single-cell level (dark blue, main text Figure 2), and burst size (light blue, main text Figure 3) models. Latent period mean, CV, and their confidence intervals were calculated as previously described. Note that the population-level inference estimates a single-valued parameter for burst size that does not account for variability. The effective burst size – defined as the average burst size of the population– and the burst size CV for the single-cell level estimate were computed using burst sizes larger than 1 for individual cells, starting at 9 hours post-infection ($n = 213$). Confidence intervals were obtained by bootstrapping the data $10^4$ times and calculating the 95% quantiles.

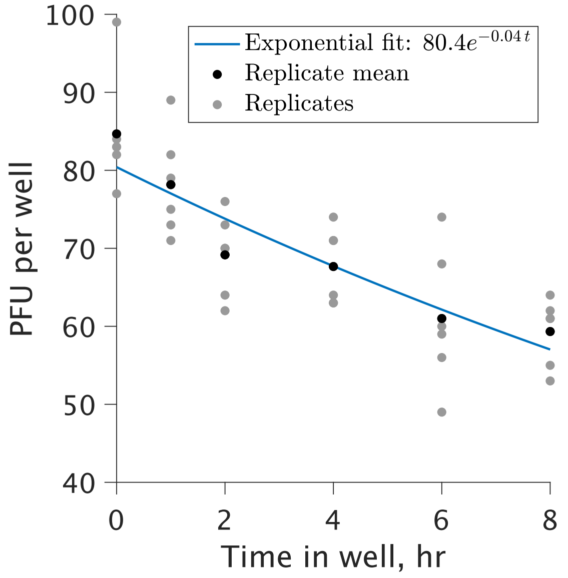

Figure S8: **Loss of viral particles due to adhesion to well plate surface**. Viral particle adhesion to the well plate surface can lead to an underestimation of individual cell burst size. We quantified this loss by adding infective viruses to 96-well plates, then periodically tested the well contents using plaque assays. The loss of infective viruses due to adhesion follows an exponential decay model (blue line) with greater losses occurring over longer periods of time in the well.

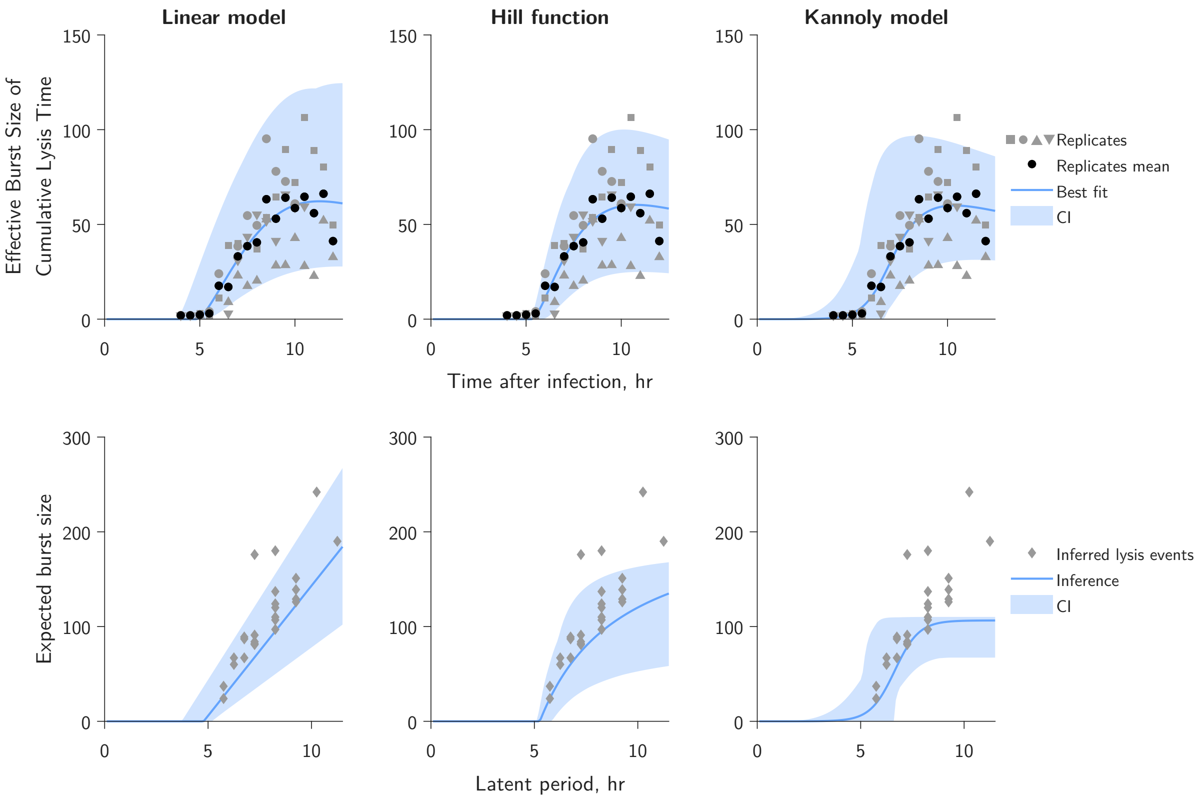

Figure S9: **Uncovering the latent period to burst size relationship**. We aimed to find the latent period to burst size relationship by fitting the effective burst size of cumulative lysis time to the entanglement of the previously predicted latent period distribution and three potential models: a linear model (left panel), a Hill function (middle panel), and a logistic growth function (right panels). Fits are shown in top panels with shaded area representing the 95 % Confidence Interval. The solid line represents the best fit. The different-shaped gray symbols indicate the different biological replicates. Black circles mark the average value over all replicates. Based on burst size distributions of sampling time points we predict the lysis events occurring at half-hour intervals (see Materials and Methods of main text). These predictions (diamonds, bottom panels) resemble a linear model. Note that the burst sizes from inferred lysis events (diamonds) were not used for inference. The solid line and shaded area show the corresponding best fit and CI in the top panel.

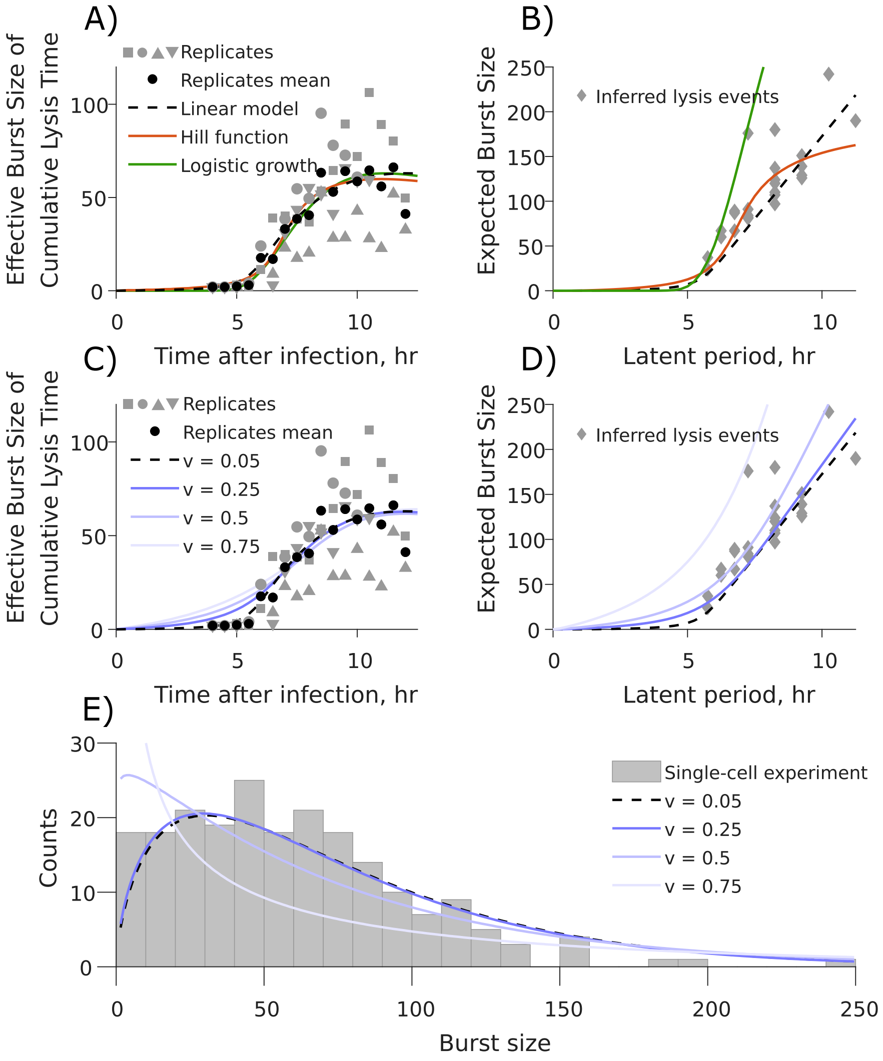

Figure S10: **Estimating the post-eclipse period to burst size relationship**. We aimed to find the latent period to burst size relationship in a scenario where the latent period is divided into eclipse and post-eclipse periods. The post-eclipse period to burst size relationship was inferred by fitting the effective burst size of cumulative lysis time to the entanglement of the previously predicted latent period distribution (Figure S3, Table S3) and three potential post-eclipse period to burst size models: a linear model, a Hill function, and a logistic growth function sourced from Kannoly et al. [8]. (A) The best fits for the linear model, Hill function, and logistic growth model are shown in dashed black, solid orange and green, respectively. We found the linear model to outperform the Hill function and logistic growth model. (B) Based on burst size distributions of sampling time points we predicted the lysis events occurring at half-hour intervals (see Materials and Methods of main text). These predictions further validate the linear model. Inferred parameters are available in Table S6. (C) Fits for the linear model for varying values of $v$, the proportion of latent period variance explained by the eclipse period. The best performing model includes a $v$ value of 0.05 (dashed line), and is the same fit shown in panel (A). (D) Expected burst size as a function of latent period predicted from the linear model with varying $v$. The diamonds correspond to predicted lysis events and were not used for parameter inference. (E) The burst size distribution expected for burst size variability deriving from latent period variability assuming the linear model for multiple values of $v$.

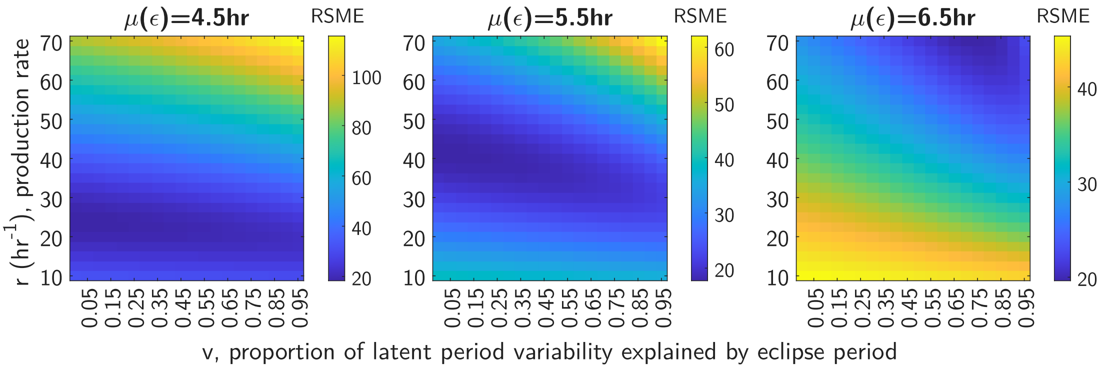

Figure S11: **The relative contribution of the eclipse period variability is non-identifiable**. Heatmaps show the Root Mean Squared Error (RMSE) between model predictions (assuming a linear post-eclipse period to burst size relationship) and data for combinations of $v$, the proportion of latent period variability explained by eclipse period variability, and $r$, the viral production rate, while the average eclipse period, $\mu(\epsilon)$, is fixed. Colors indicate the RMSE of the fit. The extended region of low RMSE values along the x-axis shows the model performance is largely insensitive to the $v$ parameter, indicating that it is non-identifiable by the data.
